# Light and temperature sensitive seizures are regulated by spatially distinct cortex glial populations in the central nervous system

**DOI:** 10.1101/2025.06.27.661959

**Authors:** Govind Kunduri, Tanja Angela Godenschwege, Katherine Sankey, Kandahalli Venkataranganayaka Abhilasha, Usha R. Acharya, Jairaj K. Acharya

## Abstract

**Significance statement:** Astrocytic glia are present throughout mammalian brain. Whether astrocytes in different parts of the brain regulate neuronal function in spatially circumscribed manner is an area of intense research. Cortex glia of *Drosophila* shares significant similarities with mammalian astrocytes in their ability to interact with neuronal soma. In this study, we established intersectional methods to restrict expression of genes of interest to cortex glia specifically located in optic lobe or central brain or ventral nerve cord. Using epilepsy models, we show that light- and temperature-sensitive seizures are differentially regulated by spatially distinct cortex glia sub-population. Central brain specific cortex glia is important for regulating light-inducible seizures, whereas ventral nerve cord specific cortex glia is essential for suppression of temperature-sensitive seizures.

Epilepsy is a brain disorder, characterized by recurrent seizures due to abnormal neuronal activity originating from a population of cortical neurons. It is known that seizures are often associated with abnormal glial cell function at the seizure focus. Recent studies have shown that each glial type such as astrocytes display significant degree of heterogeneity in their development, molecular signatures, and function depending on the brain region in which they are located. It is unknown if such heterogeneity differentially influence/cause seizures. Previous studies in *Drosophila* have shown that aberrant cortex glial function led to light inducible seizures in Ceramide phosphoethanolamine synthase (*cpes*) and temperature inducible seizures in *zyd* mutants. Here, we have optimized Gal4/Split-Gal4/Gal80/LexA drivers to specifically express a gene of interest throughout development in cortex glial subpopulations in different parts of the brain including optic lobe (OL), central brain (CB) and ventral nerve cord (VNC). Using these tools, we performed brain region specific cortex glial rescue experiments in *cpes* and *zyd* mutants. We found that OL and CB, but not VNC specific cortex glial expression of UAS CPES, were able to significantly suppress light inducible seizures in *cpes* mutants. In contrast, VNC but not OL or CB specific cortex glial expression of UAS Zyd suppressed temperature sensitive seizures. Further, in a third model, expression and activation of transient receptor potential (dTrpA1) just in the VNC specific cortex glia was sufficient to induce temperature sensitive seizures in wild type flies. Our findings suggest that regionally specialized cortex glial subtypes differentially regulate seizure susceptibility in seizure models.

## Introduction

Glia and neurons constitute the two major components of the nervous system across both vertebrates and invertebrates. Although discovered in 1854, much of our understanding of glia as regulators of neuronal differentiation, organization, function and survival has come about in the last two or three decades (1, 2). Recent literature highlights the shift in emphasis on viewing them as cohabitants and equal partners in the functions of the central nervous system rather than as scavengers and helpers of the neurons (2–5). Consequently, central nervous system pathologies are also being viewed as primary defects in either neurons or glia and not as being mainly a disease of the former (6). A classic example of this changing notion is seen in studies on epilepsy (7). Epilepsy is a central nervous system (CNS) disorder characterized by recurrent seizures due to high frequency action potentials and hyper-synchronization of a population of cortical neurons. It is estimated that epilepsy affects over 65 million people worldwide and approximately 1 in every 100 people experience seizures in their lifetime (8, 9). Development of seizures in epilepsy in humans and experimental models is often correlated with complex structural, and functional changes in astrocytic glia (reactive astrogliosis) in seizure focus that arise from CNS insults and pathology (9). Astrogliosis was considered more reactive, secondary to successive ictal insults from defective neurons that caused inflammatory, reparative scarring in the glia that could further confound the epileptic pathology. Recent epileptology studies now include reports of primary astrogliosis as a driver of epilepsy along the same vein as a major neuronal defect (10, 11). One of the first examples of this is Alexander disease, where variants of GFAP that are largely produced in astrocytes cause epileptic seizures as one of the disease’s primary symptoms (12, 13).

Understanding where in the brain abnormal neuronal activity begins and how it spreads to rest of the brain to induce seizure remains a critical goal in epilepsy research. Seizure focus can be a small region of the brain from which abnormal neuronal activity (seizure) originate. Seizure initiation site/or seizure focus have been described to be less than 1cm in diameter (14). It has been suggested that acute focal seizures begin as local synchronization of neuronal ensembles within initiation site which then spread to larger areas in a saltatory fashion to break into neighboring cortex where it proceeds smoothly to the rest of the brain (15). Although anti-epileptic drugs targeting neuronal mechanisms alleviate if not completely remove seizures in 70% of the patients, 30% of the patients remain resistant to these drugs suggesting an urgent need for investigating non neuronal mechanisms in epilepsy (16). As indicated earlier, growing number of studies have shown that glial cells including astrocytes, microglia and oligodendrocytes play crucial roles in epilepsy and thus they could be explored as potential therapeutic targets (16). Glial cells are described based on their functions and their ontogeny. Astrocytes are one of the major glial cells found in the vertebrate brains. They tile the central nervous system (in a non-overlapping manner) and are important modulators of neuronal activity (17). Different parts of the brain are populated by highly specific neurons and not surprisingly astrocytes do show structural, and molecular heterogeneity. *In vitro*, glia isolated from different regions show differences in their ability to differentiate adult rat hippocampal neural stem cells into neurons. Studies continue to parse out the regional specificities of glial networks in specific neuronal functions and patterns of activity. Historically, glial cells were studied as homogenous group of cells across brain regions (18, 19). For instance, all astrocytes across the brain regions were thought to be structurally and functionally identical (18, 19). Recent studies have uncovered a surprising degree of regional heterogeneity in their development, molecular signatures, and physiology. However, little is known about the functional significance of such regional heterogeneity of glial cells (18–20). It was suggested that glial heterogeneity across brain regions enables development or optimization of local neuronal circuit functions (18, 19). There is a distinct possibility for existence of functional diversity among the same group of glia across different regions of the brain. Elucidation of such functional specialization would allow for better understanding of physiology and management of pathology of the nervous system. Understanding heterogeneity in glial cell properties across brain regions and their dysfunction in the context of epilepsy is an outstanding question in the field that needs to be addressed for developing targeted treatments (9).

Many of our insights into glial physiology and defects that lead to human disorders have come from both vertebrate and invertebrate animal models (3). Possibility of glial heterogeneity within the same class, across brain regions have also been alluded to in a genetically tractable model organism *Drosophila melanogaster* (21–23). For instance, it was shown that different enhancers drive expression of Gal4 in specific subsets of cortex glia located in different parts of the brain including optic lobes (OL) central brain (CB), and ventral nerve cord (VNC) (22).

However, the functional significance of such regional heterogeneity in cortex glia remains unknown. Cortex glia is also known as neuronal cell body associated glia that appears as a honeycomb-like structure in the neuronal cortex, wherein each cortex glial cell extends its membranes to envelop up to 100 neuronal cell bodies (21, 24–26). Unlike mammalian astrocytes which simultaneously interact with neuronal cell bodies and synapses, cortex glia in *Drosophila* makes contacts only with neuronal cell bodies and secondary axon tracks but not with synapses. Recent studies have shown that cortex glia play important roles in regulation of neuronal hyperexcitability (27–29). Previously we have shown that ceramide phosphoethanolamine synthase (*CPES*) plays an important role in membrane expansion, and absence of CPES leads to defective cortex glial plasma membrane and development of light inducible seizures (27, 30). We have shown that altered glial plasma membrane structure, particularly failure to establish detergent resistant membranes (DRMs) is responsible for cortex glial encapsulation defect in *cpes* mutants. It has been shown that loss of function of cortex glial specific Na^+^/Ca+, K^+^ exchanger (*zyd*eco), one of the first identified glial specific gene involved in epileptic phenotype, results in temperature sensitive seizures. Zydeco (*zyd^1^*) has been shown to influence glial Ca^2+^ signaling *via* regulation of microdomain Ca^2+^ oscillations at the plasma membrane. The *zyd^1^*mutant cortex glia show higher intracellular Ca^2+^ levels (31). Further, it has been shown that increased cortex glial Ca^2+^ levels cause hyperactivation of calcineurin-dependent endocytosis of K2P channel, Sandman resulting in the reduced ability of glial cells to buffer K+ around neuronal cell bodies resulting in increased neuronal excitability (29). In this study, we optimized methods to express a gene of interest in cortex glial subpopulations in different parts of the brain including OL, CB and VNC. Subsequently, using these tools and cortex glial subpopulation specific rescue experiments, we show that OL and CB specific cortex glia are important in regulating light inducible seizures in *cpes* mutants. In contrast, VNC specific cortex glia plays an important role in regulating temperature sensitive seizures in *zyd^1^* mutants. Further, we show that Ca^2+^ influx into VNC specific cortex glia was sufficient to induce temperature sensitive seizures in wild type flies. Together our results suggest that cortex glial subpopulations across brain regions differentially regulate seizure susceptibility in different seizure models.

## Results

### Optimization of gene expression in cortex glial subpopulations located in OL, CB and VNC

To express genes of interest in a subset of cortex glial population across brain regions we started utilizing Gal4, and split-Gal4 lines generated by the Janelia Fly Light project (32–34). Gal4 driver lines that label cortex glia in different brain regions, including OL, CB, and VNC have been described before (22). It was suggested that R65B12-Gal4, R9F07-Gal4 and R54D10-Gal4 labeled the cortex glia of OL, CB, and VNC respectively in the adult brain (22). However, their precise expression patterns during developmental stages were not described. Transgenic flies corresponding to R54D10 and R9F07-Gal4 were not available at the stock centers. To obtain flies that specifically express the gene of interest in a subset of cortex glial population across brains regions, we decided to regenerate R54D10 and R9F07 transgenic flies. We first PCR amplified and cloned R54D10 and R9F07 enhancers using the primers described within the Janelia Fly Light website. We subcloned PCR products into split-Gal4 DNA binding domain (Gal4.DBD) (pBPZPGal4DBD, Addgene#26233), Gal80 (pBPGal80UW6, Addgene#26236) and nlsLexAp65 (pBPnlsLexA p65UW, Addgene #26230) vectors. Sequence verified clones were then used to generate transgenic flies as described in the methods section. The transgenic flies that were generated in this study include R54D10-Gal4.DBD, R54D10-Gal80, R54D10-nlsLexAp65, R9F07-Gal4.DBD, and R9F07-Gal80. We have combined these transgenic flies with previously published Gal4/SplitGal4 lines to promote or restrict gene expression in selective brain regions. We also browsed large Gal4/split Gal4/LexA collections at Janelia Fly Light and identified several lines including Vienna tile (VT) lines that are likely to express in cortex glia of adult central nervous system (Table 1). We chose the split-Gal4 system since Gal4 lines that are driven by single segment of genomic DNA enhancers often show expanded gene expression patterns (32, 35). On the other hand, intersectional methods such as split-Gal4, wherein the full length Gal4 transcription factor is split into DNA-binding domain (DBD) and an activation domain (AD) that can utilize two different enhancers to restrict gene expression to overlapping areas of expression. Each domain by itself is insufficient to bind to upstream activating sequence (UAS) and drive transcription. However, when both AD and DBD are present in a given cell they interact with each other *via* the leucine zipper-dimerization domain and form a functional transcription factor which in turn binds to UAS elements to activate transcription (32).

Previous studies have shown that Nrv2-Gal4 specifically labels all cortex glial subtypes and ensheathing glia in adult brain. The Nrv2-p65(AD) flies were a kind gift from JC Coutinho-Budd (21). We have used UAS mCD8GFP (to visualize the cell membrane) and UAS mCherryNLS (to visualize the cell nucleus) flies as reporters of Gal4/split Gal4 mediated gene expression. Our screen for cortex glial specific drivers at FlyLight collection identified Viena tile line VT038983 as a potential candidate that strongly expresses in cortex glia of adult CB and VNC but weakly in OL (Table 2). To restrict the expression of a gene of interest just in CB and VNC, we recombined Nrv2-p65(AD) with VT038983-Gal4.DBD and drove the expression of UAS mCD8GFP and mCherryNLS. In order to examine expression patterns of this split-Gal4, we dissected central nervous system (CNS) from 3^rd^ instar, 48 hours post pupation and 7-day old adult stages, immunostained and analyzed them by confocal microscopy as described in the methods section. As shown in Fig.1(a-c), we found that this combination specifically labeled cortex glia in all developmental stages including 3^rd^ instar, pupa and adult stages. Interestingly, it did not label OL specific cortex glia during 3^rd^ instar and pupal stages, as a result expression is primarily limited to CB and VNC during these developmental stages (Fig.1a&b; Table 2). However, in the adult brain this combination begins to express in OLs also, resulting in more global/generic cortex glial expression pattern (Fig.1c). When we combined R9F07-Gal4.DBD and VT038983-p65(AD) Fig.S1(a-c) the expression in larval and pupal stages was similar to Nrv2-p65(AD) with VT038983-Gal4.DBD. In the adult while it mostly followed the larval and pupal pattern, one of the two optic lobes showed weak expression Fig.S1(c), suggesting that this combination of split-Gal4 expression is mostly restricted to CB and VNC.

**Fig.1.**
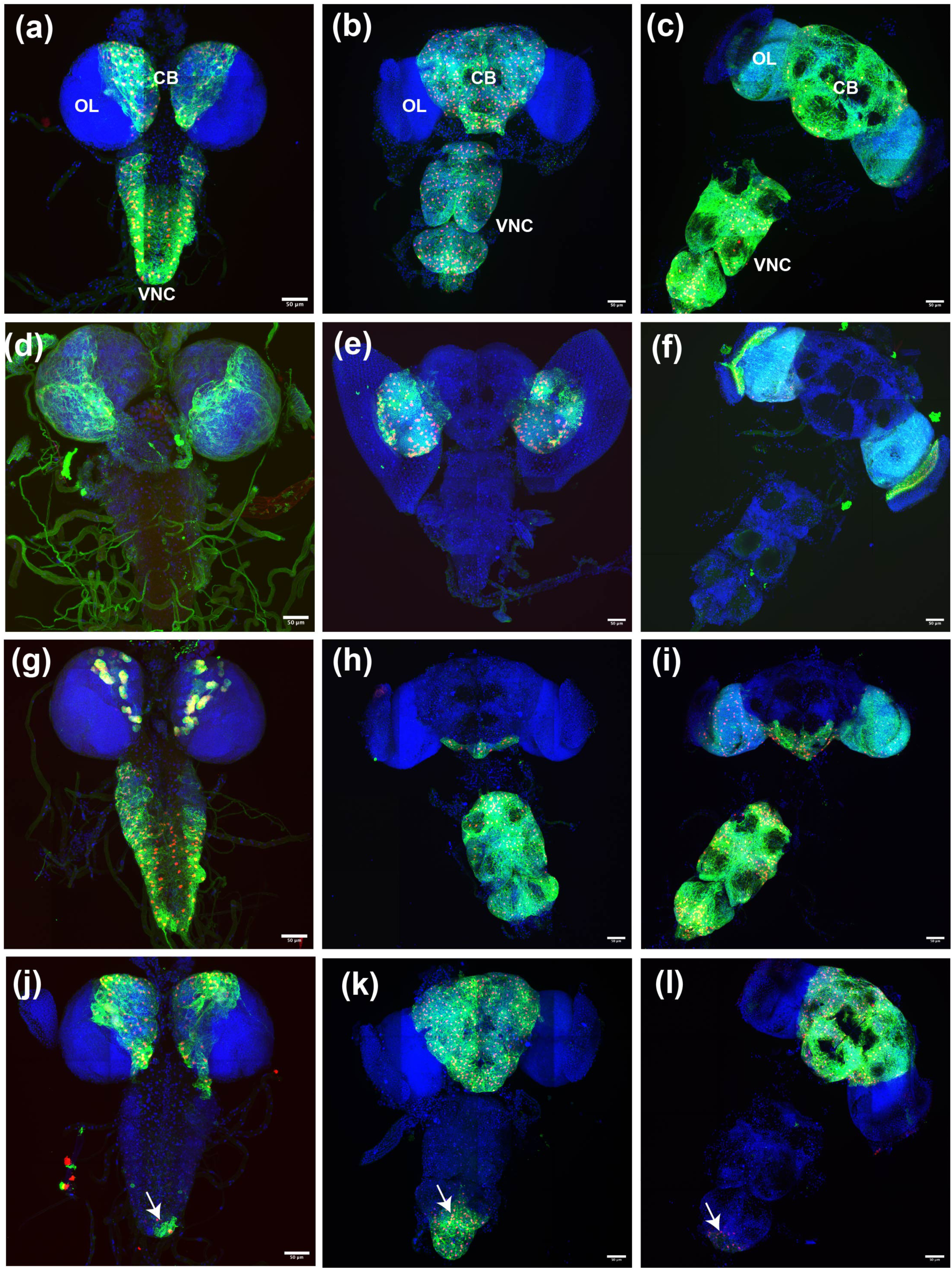
Labeling region-specific cortex glial cells in *Drosophila* central nervous system. **(a-c)** Expression pattern of Nrv2-p65 (AD), and VT038983-Gal4.DBD in fly CNS, during 3^rd^ instar larval stage (a), 48 hours post pupation (b) and 7-day old adult (c). **(d-f)** Expression pattern of R65B14-Gal4, nSyb-Gal80 and R9F07-Gal80 in fly CNS during 3^rd^ instar (d), 48 hours post pupation (e) and 7day old adult (f). (g-i) expression pattern of VT038983-p65 (AD) and R54D10-Gal4.DBD during 3^rd^ instar (g), 48hours post pupation (h) and 7-day old fly CNS (i). **(j-l)** Expression pattern observed with the combination of R54D10-nlsLexAp65, 13xLexAop2 KZip+. 3xHA and Nrv2-p65 (AD), VT038983-Gal4.DBD, during 3^rd^ instar (j), 48 hours post pupation (k) and 7-day old fly CNS (l). Arrows indicate continued expression of markers at the tip of VNC. The UAS mCD8GFP and mCherry NLS were used as reporters for the above-mentioned combination of Gal4/splitGal4/LexA/Gal80 constructs. Scale bar in each image corresponds to 50 micrometers.

R65B12-Gal4 has been shown to label OL specific cortex glia of adult brain (22). Our analysis of R65B12-Gal4 through the developmental stages showed that in addition to specific expression in the cortex glia of OLs, it is also expressed in neurons of the 3^rd^ instar brain Fig.S1(d) and is expressed in cortex glia of the CB and VNC during pupal stages Fig.S1(e). As reported previously, in the adult brain R65B12-Gal4 expression is restricted to OL specific cortex glia, and weakly in neurons Fig.S1(f). Due to lack of alternative Gal4 drivers that strictly express in cortex glia of the OLs, we decided to optimize R65B12-Gal4 expression through the development of flies. Gal4 dependent transcriptional activation of UAS transgenes is inhibited by co-expressing Gal80 repressor that binds to Gal4 activation domain. To prevent R65B12-Gal4 expression in neurons, we co-expressed it with pan neuronal Gal80 (nsyb-Gal80). Similarly, to prevent leaky expression of R65B12 in CB and VNC specific cortex glia during pupal stages, we co-expressed it with R9F07-Gal80 which in turn blocked the expression in cortex glia of CB and VNC but not in OLs. Consistently, we found that co-expression of R65B12-Gal4 with nsyb-Gal80 and R9F07-Gal80 limited its expression to optic lobe specific cortex glia throughout the developmental stages Fig.1(d-f) (Table 2).

In adult brain R54D10-Gal4 expresses specifically in cortex glia of VNC and subesophageal ganglion (SEG) (22). To examine its expression throughout development, we utilized the split-Gal4 system wherein we have combined Nrv2-p65(AD) or VT038983-p65(AD) (activation domains expressed in all cortex glia) with the R54D10-DBD (VNC specific cortex glia) to drive expression of mCD8GFP and mCherry NLS. As shown in Fig.S2 (a & b), Nrv2-p65(AD)-R54D10-Gal4.DBD expression is primarily restricted to VNC in 3^rd^ instar and pupal brains. However, in adult brains we found that in addition to the VNC, and SEG, gene expression is also visible in OLs Fig.S2 (c) (Table 2). Overall, consistent with the previously published results, R54D10 expression was completely absent in major part of the 3^rd^ instar, pupal and adult CB. We found similar results while using VT038983-p65(AD), and R54D10-Gal4.DBD Fig.1(g-i), although we have noted some additional expression in cortex glia of neuroblast colonies in the CB region of 3^rd^ instar brains Fig.1(g), this expression did not continue in pupal and adult CBs Fig.1(h & i). Together, these results suggest that R54D10 expression is mostly excluded in the CB and the expression is primarily restricted to VNC, SEG and OL.

Currently there are no Gal4/split-Gal4 drivers available to express a gene of interest specifically in cortex glia of the CB. To achieve CB specific cortex glial expression, we have combined split-Gal4 system with the LexA Killer zipper (KZip+) system. In this method, a dominant negative repressor (Gal4DBD/KZip+) is expressed under the control of a third promoter, which in turn represses the split-Gal4 mediated gene expression in a subset of cells where it is expressed (36). As shown in Fig.1(a-c), the split-Gal4 (Nrv2-p65(AD)-VT038983-Gal4.DBD) was expressed in cortex glia of almost all brain regions in adult brain. To limit its expression only to CB we decided to repress gene expression in VNC and OL of this split-Gal4 using the killer zipper approach. To this end, we have expressed nlsLexAp65 under the control of R54D10 enhancer, which we have shown to be expressed primarily in VNC, SEG, and OL of adult brain Fig.1(g-i) and Fig.S2(a-c). The nlsLexAp65 in turn binds to 13x LexAop2 promoter to produce KZip+ in VNC, SEG, and OL Fig.S3(d-f), thereby leaving expression only to CB. Consistently, as shown in Fig. 1 (l), in adult brains, the gene expression is primarily restricted to CB. However, repression was not complete at the tip of the VNC in 3^rd^ instar, and pupal brains, perhaps due to insufficient production of KZip+ in that area during development Fig.1 (j &k). Adult brains however showed a stronger repression, as a result only a weak expression is visible at the tip of VNC Fig.1 (l). We found similar results with another split-Gal4 combination including Nrv2-p65(AD)-Wrapper-Gal4.DBD (Ctx-Gal4) Fig.S3(g-i). Taken together R54D10 Killer zipper successfully represses split Gal4 mediated gene expression in most part of the VNC and OL during development, thereby leaving expression mostly to CB (Table 2).

### Central brain specific cortex glia is important in light inducible seizures

Having largely optimized Gal4/split Gal4 drivers that express a gene of interest in a subset of cortex glia across brain regions across developmental and adult stages we asked whether cortex glia in different brain regions regulate seizures differently in different seizure models such as light and temperature inducible seizures. We have previously shown that ceramide phosphoethanolamine synthase (CPES) null mutants display light inducible seizures. In *cpes* mutants, cortex glial plasma membranes fail to envelop neuronal cell bodies resulting in loss of glial cell support to neurons leading to development of light inducible seizures (27). To identify brain region that play important role in light inducible seizures, we have performed brain region specific cortex glial rescue experiments using optimized Gal4/SplitGal4/LexA drivers expressing UAS CPES. We hypothesized that UAS CPES expression in one part of the brain should rescue the cortex glia in the respective region. For instance, expression of UAS CPES in the VNC should rescue the cortex glia in the VNC only but not in the optic lobes or central brain. Similarly, expression of UAS CPES in the OL or CB should rescue the cortex glia only in the OL or CB respectively. To investigate, if this is the case, we have first labeled cortex glia using generic cortex glial driver Wrapper-nlsLexAp65 (R54H02-nlsLexAp65) and membrane specific protein LexAop2-rCD2-GFP. As shown in Fig.2 (a&b), compared to control Fig.2a, *cpes* mutant brains Fig.2b show defective cortex glia in all parts of the brain including VNC, CB, and OL.

**Fig.2.**
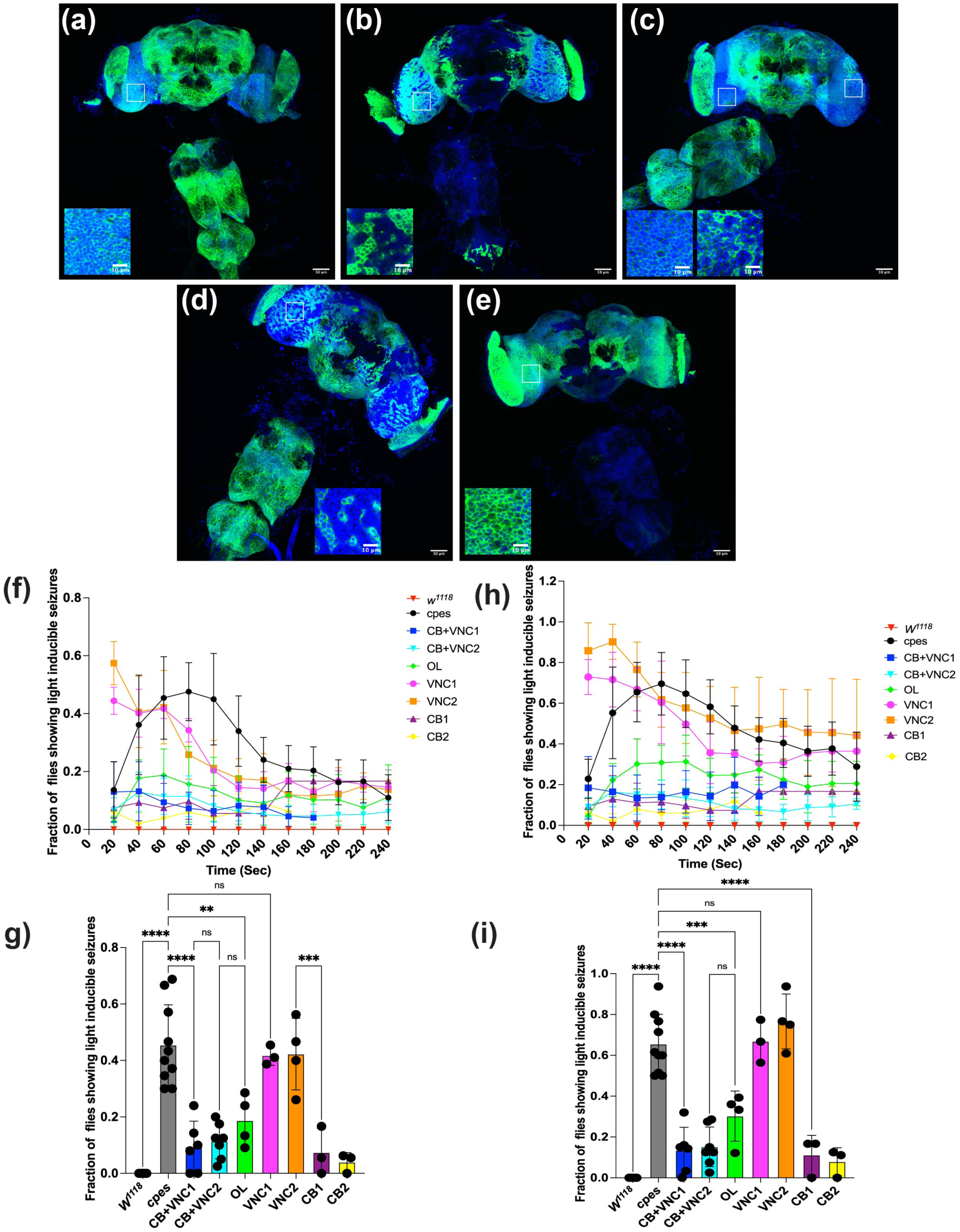
Evaluation of defective cortex glial and light inducible seizure rescue with region-specific Gal4 drivers in *cpes* mutants. (a-e) Adult CNS, wherein cortex glia is labeled with Wrapper-nlsLexA-p65 and LexAop2-rCD2-GFP, in various genetic backgrounds including in wild type (a); *cpes* mutant with UAS CPES alone (b); *cpes* mutant background with UAS CPES expressed via split Gal4, Nrv2-p65 (AD), and VT038983-Gal4.DBD (c); *cpes* mutant background with UAS CPES expressed in the ventral nerve cord with the split Gal4 combination VT038983-p65(AD) and R54D10.Gal4.DBD (d). *cpes* mutant background with UAS CPES expressed in the optic lobes with the combination of R65B12-Gal4, nSyb-Gal80, R9F07-Gal80 (e); Inset shows a small portion of optic lobe specific cortex glia in all backgrounds (a-e). (f-g) quantification of fraction of flies showing complete seizures (i.e., flies fell to the bottom and lying on their back or side) as a function of time. (g) statistical analysis for the complete light inducible seizures at time point 1min/60 sec. (h-i) Quantification of fraction of flies showing complete and partial seizures (i.e., flies fell to the bottom and remain standing without movement for at least 20 sec). (i) Statistical analysis for the complete and partial light inducible seizures phenotype at time point of 1 min/60 sec. Each data point in (f) and (h) represent a minimum of 50 flies from 3 independent experiments. Each dot in (g) and (I) represents an independent experiment consisting of 15-30 flies. CB+VNC1=Nrv2-p65(AD), VT038983-Gal4.DBD>UAS CPES; CB+VNC2= VT038983-p65(AD) and R9F07-Gal4.DBD>UAS CPES; OL=nSybGal80, R9F07Gal80, R65B12-Gal4>UAS CPES; VNC1= VT038983-p65(AD), R54D10-Gal4.DBD>UAS CPES; VNC2=Nrv2-p65(AD), and R54D10-Gal4.DBD>UAS CPES; CB1=Nrv2-p65(AD),VT038983-Gal4.DBD, 13xLexAOP2 Kzip+, R54D10-nlsLexAp65 > UAS CPES; CB2= Nrv2-p65(AD), Wrapper-Gal4.DBD, 13xLexAOP2 Kzip+, R54D10-nlsLexAp65> UAS CPES; Central brain (CB), ventral nerve cord (VNC), and optic lobe (OL). Scale bar in images (a-e) corresponds to 50 micrometers and 10 micrometers for the inset images.

Expression of UAS CPES in CB and VNC specific cortex glia (Nrv2-p65(AD) and pVT038983-Gal4.DBD) in *cpes* mutant background, fully restored the cortex glial defects in respective brain regions. However, we observed cortex glia in OL is also partially rescued (compare left and right insets in Fig.2c). Interestingly, expression of UAS CPES in VNC specific split-Gal4 (VT038983-p65(AD), and R54D10-Gal4.DBD) restored cortex glia in VNC and SEG, but not in the CB and OL, consistent with its expression pattern during development (compare Fig.2d with the Fig.1g-i). Although pVT038983-p65(AD), R54D10-Gal4.DBD also drives weak expression in the OL of the adult brain (Fig.1i), no rescue of cortex glia was observed in this region (Fig.2d), likely due to its late expression during development. That is, presence of CPES during larval and pupal stages is essential for rescue of cortex glia, and late expression in adult brain may not restore cortex glia that was already disrupted during development. Similarly, we found that expression of UAS CPES in OL specific cortex glia (R65B12-Gal4, R9F07Gal80) fully restored cortex glia in OL, but not in the VNC region, consistent with the expression pattern (Fig.2e). We also observed partial rescue in CB specific cortex glia, in addition to OL specific rescue (compare Fig.2b and Fig.2e), probably due to insufficient repression of R65B12-Gal4 expression in the CB by R9F07-Gal80 during pupal stages (Fig.S1d-f). However, it should be noted that, unlike mCD8GFP and mCherryNLS expression, CPES expression is an extremely sensitive reporter of gene expression, since each enzyme molecule is likely to produce thousands of lipid molecules which may easily rescue cortex glial defects. Taken together these results suggest that brain subregion specific cortex glial rescue with UAS CPES mostly matches with the expression pattern observed for the corresponding Gal4/split-Gal4 drivers with fluorescent reporters.

To further investigate how brain subregion specific cortex glial rescue affects light inducible seizures, we have expressed UAS CPES using one OL, two sets of CB, and two sets of VNC specific Gal4/Split Gal4 drivers. The adult CNS region-specific rescue flies in the *cpes* mutant background were collected and aged for 3-4 weeks and measured for light inducible seizures as described in the methods section. The *cpes* mutant adult flies are sensitive to fluctuations in light intensity and show robust seizures when shifted from dark to ambient light condition (27). They typically show two behavioral phenotypes upon shifting from dark to light, including, falling to the bottom and lying on their back or sides until recovery (complete seizure/paralysis) or falling to the bottom and remaining immobile while standing on their legs (partial seizure). As shown in Fig.2f&g (complete seizures) and Fig.2h&i (complete and partial), *cpes* mutant flies showed significant amount of light inducible seizures compared to wild type flies. Fig.2g and Fig.2i represent statistical analysis for the data in Fig.2f and Fig.2h respectively at 60 seconds time point. Expression of UAS CPES in the cortex glia of CB and VNC using two different split Gal4 drivers significantly suppressed light inducible seizures, suggesting the importance of this cortex glia in light inducible seizures (Fig.2 g&i; Movie.1). Interestingly, expression of UAS CPES just in the VNC and SEG using two different split Gal4 drivers did not rescue light inducible seizures, suggesting cortex glia in the VNC regions may not play a major role in light inducible seizures (Fig.2g&i; Movie.1). It should be noted that VNC+SEG specific cortex glial rescues showed increased sensitivity to light at earlier time points and subsequently showed faster recovery kinetics compare to *cpes* mutants (Fig.2f &h; Movie.1). These results suggest that VNC+SEG specific cortex glia may not have a direct role in suppression of light inducible seizures, however they could indirectly modulate the duration of the seizure. Expression of UAS CPES using two different CB specific cortex glia drivers significantly suppressed light induced seizures (Fig.2g&i; Movie.1), suggesting an important role for CB specific cortex glia in light inducible seizures. We also observed a partial suppression of light inducible seizures with OL specific cortex glia (Fig.2g&i; Movie.1). It should be noted that as mentioned above OL specific expression of UAS CPES also partially rescues the CB specific cortex glia, suggesting this partial rescue may be due to rescue of cortex glia in the CB, which may be independent of its rescue in the OL. Taken together these results suggest that CB specific cortex glia is important in light inducible seizures of *cpes* mutants.

### Ventral nerve cord specific cortex glia is important in temperature sensitive seizures

Hypomorphic mutants of cortex glial specific Na^+^/Ca^2+^, K^+^ exchanger (*zyd*eco) suffer from strong temperature sensitive seizures (31). Zydeco (Zyd) was shown to be expressed in all cortex glia of the CNS. Unlike *cpes* mutant, *zyd^1^* mutants do not show any defects in cortex glial morphology, and acute expression of Zyd in the cortex glia was sufficient to rescue temperature sensitive seizures in *zyd^1^*mutants (31). Having identified that CB specific cortex glia are involved in the regulation of light inducible seizures, we wondered if CB region also influences temperature sensitive seizures. To investigate brain subdomains that may be involved in temperature sensitive seizures, we performed brain region specific rescue experiments on *zyd^1^* mutants using UAS-Zyd and the tools that we have optimized in this study (Fig.1). We crossed *zyd^1^*;+/+;UAS-Zyd female flies with various cortex glial specific Gal4/split Gal4 driver males and collected male progeny for measurement of heat inducible seizures. We have collected 10-20 male flies per vial and aged for 3-5 days before performing the temperature sensitive assay by exposing them to 38.5°C in a water bath as described previously (29, 31). As shown in Fig.3a&b expression of Zyd in all cortex glia completely rescued the temperature sensitive seizures in *zyd^1^* mutants. Interestingly, we find that expression of Zyd in the VNC+SEG+OL alone was sufficient to rescue temperature sensitive seizures in *zyd^1^*mutants (Movie.2). However, expression of Zyd in the CB or OL alone was not sufficient to rescue heat inducible seizures in *zyd^1^*mutants (Fig.3a &b; Movie.2). These results indicates that VNC+SEG specific cortex glia play an important role in the generation of heat inducible seizures. Although, CB or OL specific cortex glial rescue did not suppress heat inducible seizures, they did recover faster compared to *zyd^1^* mutants alone, suggesting an ancillary role for cortex glia in these regions in seizure recovery (Fig.3c&d).

**Fig.3.**
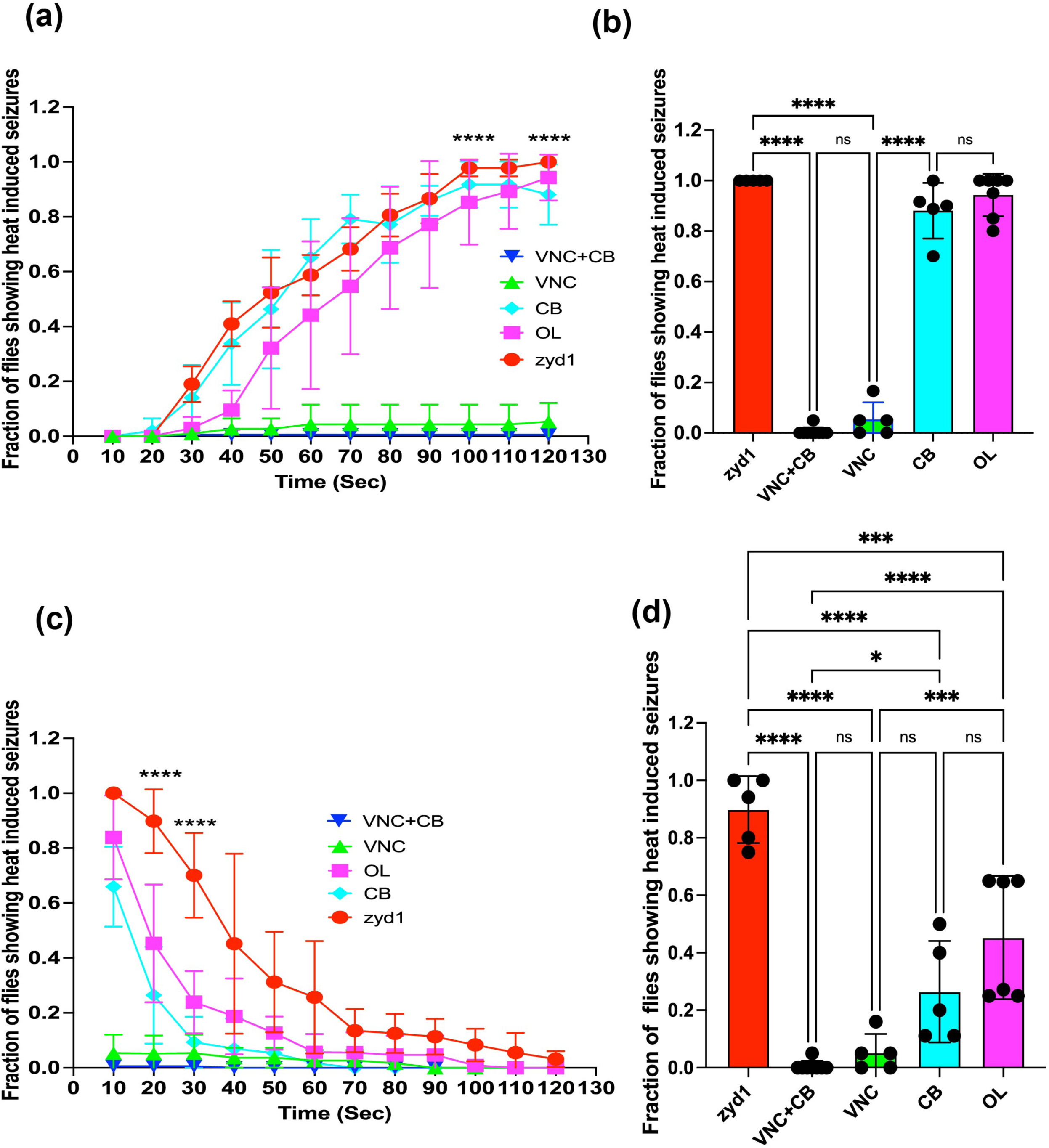
NCKX expression in the VNC specific cortex glia is sufficient for suppression of heat induced seizures in *zyd*^1^ mutants. **(a)** Measurement of heat inducible seizures in *zyd^1^* mutants and various brain region specific cortex glial rescue and its effect on suppression of heat inducible seizures as a function of time. UAS-NCKX was expressed in both central brain and ventral nerve cord (CB+VNC), ventral nerve cord alone (VNC), central brain alone (CB), optic lobe alone (OL). (b) statistical analysis of data in (a) at 120 seconds time point. (c) Measurement of seizure recovery in *zyd*^1^ mutants and brain regions specific cortex glial rescues with UAS-NCKX expression as a function to time. (d) statistical analysis of seizure recovery of data in (c) at time points of 20 seconds. Each data point in (a) and (c) represents a minimum of 50 flies from 5 independent experiments. Each dot in the (b) and (d) represent an independent experiment. VNC=VT038983-p65(AD), R54D10-Gal4.DBD> UAS NCKX; OL=R65B12-Gal4, R9F07-Gal80> UAS NCKX; VNC+CB=Nrv2-p65(AD), VT038983-Gal4.DBD>UAS NCKX.

To further investigate the importance of VNC specific cortex glia in suppression of temperature sensitive seizures in *zyd^1^* mutants, we have utilized the Gal80 system to systematically deplete Zyd expression in VNC alone and VNC+ CB (Fig.4). Previous studies have shown that Wrapper-Gal4 specifically labels cortex glia in all brain regions throughout development (Fig.4a-c & (22)). To repress wrapper-Gal4 expression in VNC, we have co-expressed it with R54D10-Gal80. Consistent with its expression pattern (Fig.1g-i & Fig.S2 a-c), R54D10-Gal80 repressed wrapper-Gal4 expression in VNC, SEG and OL in adult brain, thereby limiting membrane CD8GFP expression largely to CB (Fig.4 d-f). However, we believe R54D10 Gal80 repression is not complete in the OL of the adult brain as judged by weak detection of mCherryNLS (Fig.4f). Further, we have co-expressed Wrapper-Gal4 with R9F07-Gal80 and found that it repressed Wrapper-Gal4 expression both in the VNC and CB in all developmental stages thereby limiting mCD8GFP and mCherry NLS expression largely to OL (Fig.4g-i). Wrapper-Gal4 mediated expression of UAS-Zyd completely rescued the temperature sensitive seizures in *zyd^1^*mutants (Fig.4j). Interestingly, repression of Wrapper-Gal4 mediated UAS-Zyd expression in the VNC using R54D10-Gal80 triggered heat inducible seizures (Fig.4j; Movie.3), suggesting the importance of Zyd in VNC for suppression of seizures. Similar results were observed upon depletion of Wrapper-Gal4 mediated UAS-Zyd expression in VNC and CB using R9F07-Gal80 (Fig.4j; Movie.3). Consistent with Fig.3d, recovery analysis in these genetic backgrounds again showed that UAS-Zyd expressed in the OL alone or CB+OL showed faster recovery compared to *zyd^1^* mutants alone suggesting some role of Zyd in these regions in seizure recovery (Fig.4k). Taken together these results suggest that Zyd plays a major role in VNC but not in CB or OL to suppress temperature sensitive seizures.

**Fig.4.**
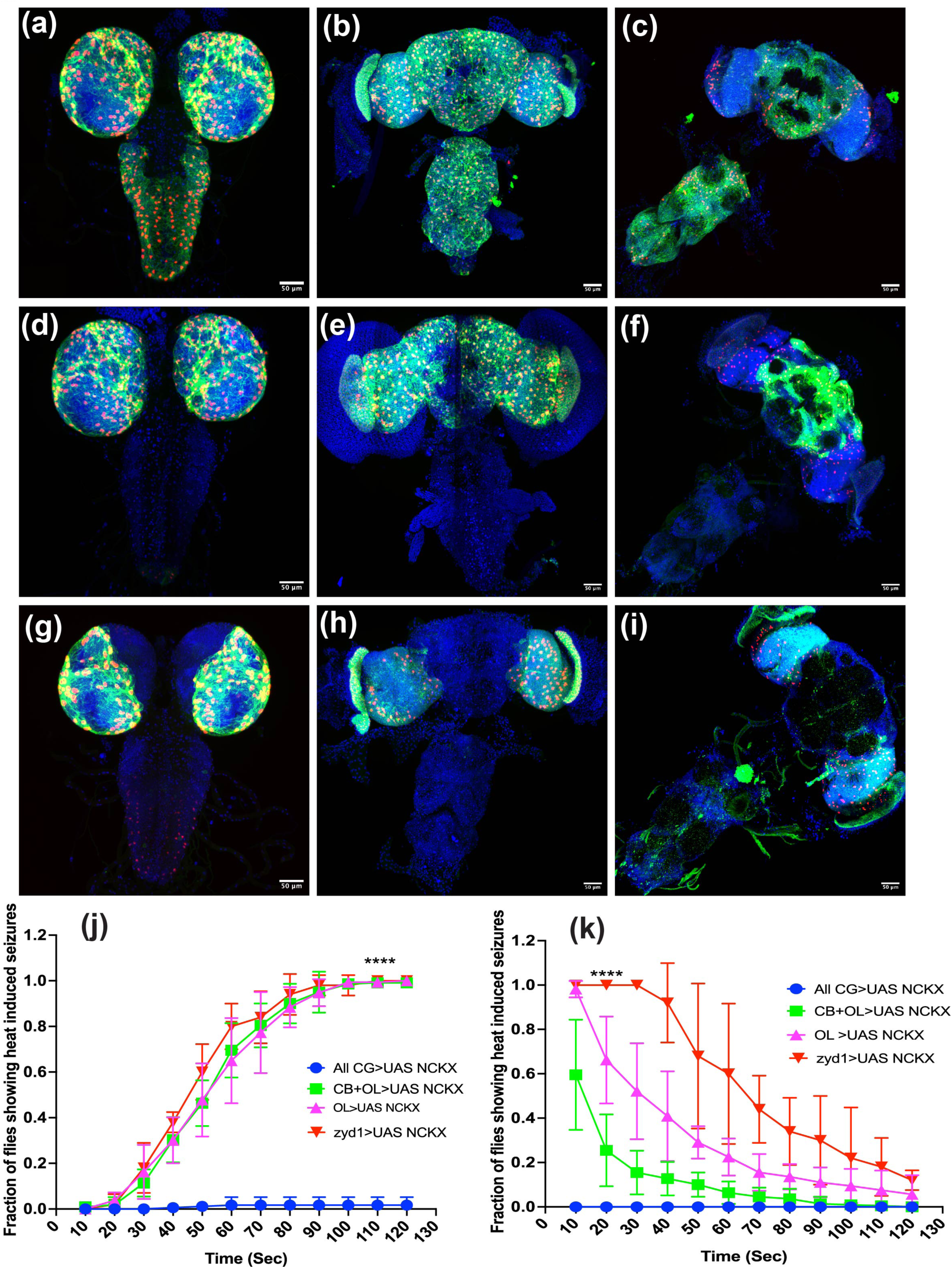
NCKX expression in the VNC specific cortex glia is essential for suppression of heat induced seizures in *zyd^1^* mutants. (a-c) Expression pattern of Wrapper-Gal4 in all cortex glia (R54H02-Gal4>mCD8GFP, mCherryNLS) in 3^rd^ instar brain (a), 48 hours post pupation (b), and 7-day old adult CNS. (d-f) Expression pattern of R54H02-Gal4 after repression with R54D10-Gal80> mCD8GFP, mCherryNLS in 3^rd^ instar brain (d), 48 hours post pupation CNS (e) and 7-day old adult CNS (f). (g-i) Expression pattern of R54H02-Gal4 after repression with R9F07-Gal80>mCD8GFP, mCherryNLS in 3^rd^ instar brain (g), 48 hours post pupation CNS (h) and 7-day old adult CNS (i). Scale bar in images (a-i) corresponds to 50 micrometers. (J) Quantification of heat inducible seizures in different genetic backgrounds, including all cortex glial rescue with R54H02-Gal4 and UAS NCKX and subsequent repression of UAS NCKX expression in VNC alone and VNC+CB using R54D10-Gal80 and R9F07-Gal80 respectively. (k) Quantification of heat induced seizure recovery of flies in (j), as a function of time. Each data point in (j) and (k) represents about 100 flies from a minimum of 5 independent experiments. All cortex glia =R54H02 Gal4>UAS NCKX; CB+OL=R54H02-Gal4, R54D10-Gal80>UAS NCKX; OL cortex glia = R54H02-Gal4, R9F07Gal80> UAS NCKX.

### Acute Ca^2+^ influx in VNC specific cortex glia via TRPA1 activation triggers heat induced seizures in control flies

Acute elevation of intracellular Ca^2+^ in cortex glia, by over expression of transient receptor potential (dTRPA1) triggers heat inducible seizures in flies (31). TRPA1 is a heat activated cation channel that is normally expressed in a subset of thermosensitive neurons (37, 38). However, when expressed ectopically it promotes Ca^2+^ influx at temperatures above 26°C but remains inactive at lower temperatures (37–39), suggesting dTRPA1 is a useful tool to acutely modulate intracellular calcium levels *in vivo* that lead to heat inducible seizures. Based on our findings in *zyd^1^* mutants, we wondered if expression of dTRPA1 in the VNC specific cortex glia would be sufficient to induce temperature sensitive seizures in a wild-type background. To test this hypothesis, we overexpressed UAS-dTRPA1 using two different sets of VNC specific cortex glial drivers (Nrv2-p65(AD)-R54D10-Gal4.DBD and VT038983-p65(AD)-R54D10-Gal4.DBD) and 3–5-day old adult flies were subjected to higher temperature. Interestingly, we found that expression of dTRPA1 in VNC+SEG+OL alone was sufficient to strongly induce temperature sensitive seizures (Fig.5a&b; Movie.4). Nrv2-p65(AD)-R54D10-Gal4.DBD mediated expression of dTRPA1 in the VNC shows stronger seizure phenotype compared to VT038983-p65(AD)-R54D10-Gal4.DBD, likely due to its stronger expression and or its expression in other types of glia such as ensheathing glia (Fig.5a&b; Movie.4). To further validate these findings, we have expressed dTRPA1 in all cortex glia using wrapper-Gal4, and subsequently repressed its expression in VNC and VNC+CB using R54D10-Gal80 and R9F07-Gal80 as described in the previous section. As expected, expression of dTRPA1 in all cortex glia strongly induced seizures at higher temperatures (Fig.5a&b). Repression of dTRPA1 expression in the VNC+SEG+OL significantly suppressed temperature sensitive seizures, while repression of dTRPA1 both in VNC and CB but not in OL completely suppressed temperature induced seizures (Fig.5a&b; Movie.4). Recovery analysis showed that dTRPA1 expressed in all cortex glia, or VNC+SEG+OL take longer time compared to when dTRPA1 expression was repressed in VNC+SEG+OL (Fig.5c&d). Taken together these results suggest that VNC specific cortex glia is essential in regulation of temperature induced seizures.

**Fig.5.**
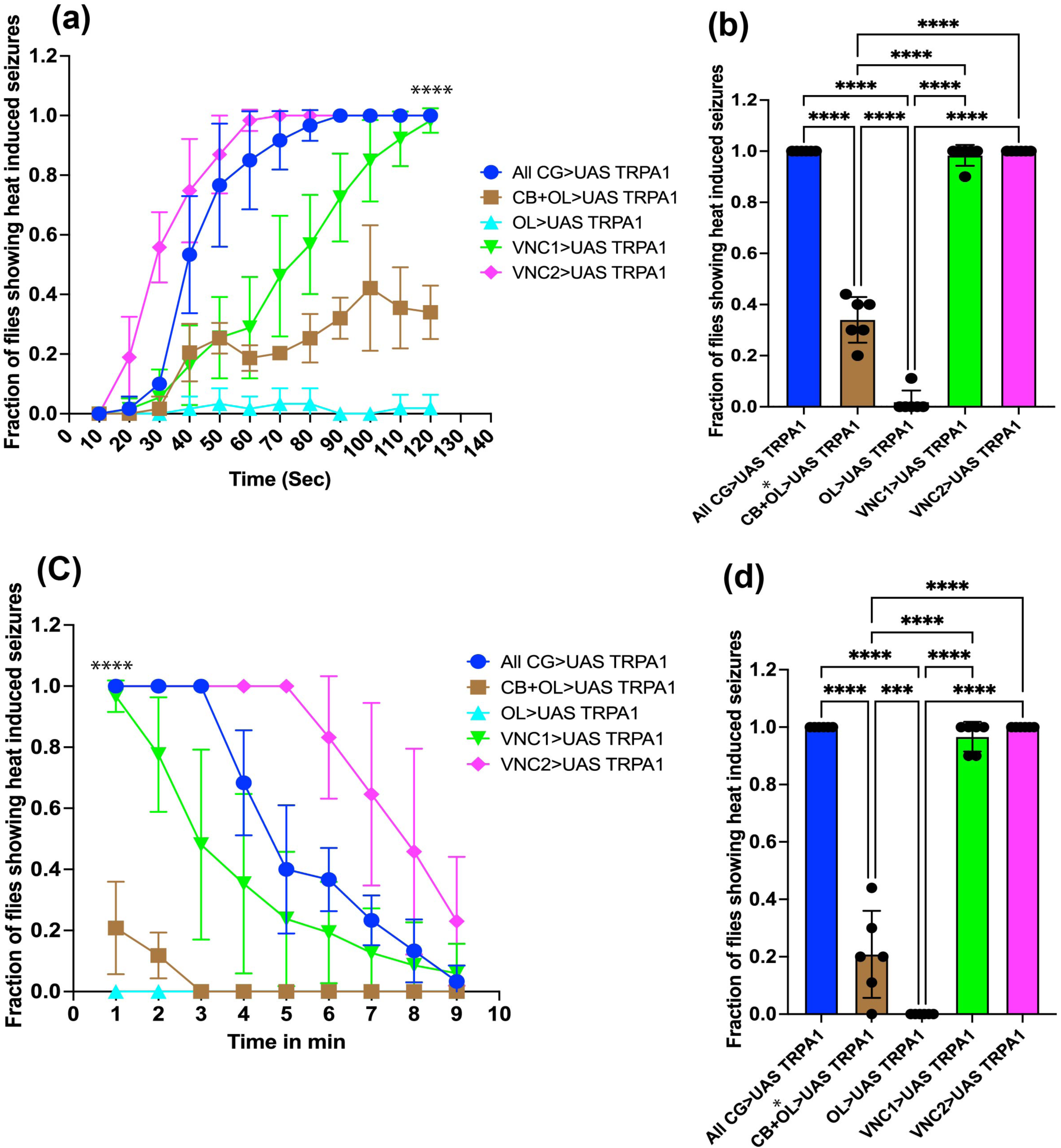
TRPA1 expression in the VNC specific cortex glia is sufficient to cause heat inducible seizures in wild type background. **(a)** Measurement of heat inducible seizures in wild type background when UAS TRPA1 is overexpressed in brain region specific cortex glia as a function of time. (b) Statistical analysis for the data in (a) at 120 seconds time point. (c) Quantification of heat induced seizure recovery as a function of time in minutes (d) Statistical analysis for the data in (c) at 1 min time point. Each data point in (a) and (c) represents a minimum of 50 flies from 6 independent experiments. Each dot in (b) and (d) represents an independent experiment. All cortex glia=R54H02-Gal4>UAS TRPA1; CB+OL= R54H02-Gal4, R54D10-Gal80>UAS TRPA1; OL=R54H02-Gal4, R9F07Gal80>UAS TRPA1; VNC1= VT038983-p65(AD), R54D10-Gal4.DBD>UAS TRPA1; VNC2= Nrv2-p65(AD), R54D10-Gal4 DBD> UAS TRPA1.

### Central brain function is dispensable for initiation of heat induced seizures

Adult *Drosophila* sense warm temperatures via internal anterior cell (AC) neurons located inside the head and peripheral hot cell (HC) neurons located in the aristae. AC and HC neurons respond to innocuous temperatures (above 25 °C) via thermoreceptors TRPA1 and Gustatory Receptor (Gr28b (D)) respectively. Both these neurons project into distinct regions of the central brain (40). Unlike adult flies, larvae respond to high temperatures via multiple dendritic (MD) neurons located in their body wall via two thermoreceptor TRP channels, including TRPA1 and Painless. TRPA1 and Painless respond to noxious warmth at temperatures above 39 °C. Although, MD neurons are also found in adult abdomen (41), it is currently unknown if they have role in noxious warmth sensing. Nevertheless, our data from temperature sensitive *zyd* mutants, and dTRPA1 overexpression analyses indicates that neuron glia interactions within the VNC+SEG region are critical for modulation of temperature induced seizures. However, it remains unclear how higher temperatures modulate neural circuits in the VNC. One possibility is that AC and HC neurons first sense higher temperatures and transmit action potentials to VNC via activation of neural circuits in the CB. Another possibility is that MD neurons located in the abdomen sense high temperatures via TRP thermoreceptors TRPA1 or Painless and transmit actional potentials to VNC. To investigate whether the central brain function is necessary for the initiation of temperature sensitive seizures, we first ablated the antenna in *zyd^1^* mutants and performed temperature sensitive assay. As shown in Movie.5 & Fig.S4a, ablation of antenna did not prevent seizure occurrence in *zyd^1^* mutants suggesting peripheral HC neurons located in the aristae are dispensable for initiation of temperature sensitive seizures. To further investigate importance of central brain function we have performed temperature sensitive assay with decapitated *zyd^1^*mutants. Interestingly, as shown in Movie.6 & Fig.S4a, headless *zyd^1^*flies continue to show seizure like behavior suggesting initiation of temperature sensitive seizures does not depend on central brain function. Similar results were observed with decapitated flies overexpressing UAS-dTRPA1 in cortex glia (Movie.6, & Fig.S4a). To investigate existence of dTRPA1, Gr28b and Painless thermoreceptor expressing neurons in VNC, we have expressed mCD8GFP and mCherry NLS using dTRPA1, Gr28b and Painless Gal4 drivers. Interestingly, we found that several of these Gal4 drivers express in VNC specific neurons in addition to CB neurons (Fig.S4b-g), suggesting their possible role in initiation of temperature sensitive seizures. To probe whether thermoreceptors dTRPA1 are directly involved in initiation of temperature sensitive seizures we have generated *zyd^1^* and *trpA^1^* double mutants and performed the temperature sensitive assay. However, no suppression of seizures was observed in *zyd^1^* and *trpA^1^* double mutants (Movie.7 & Fig.S4i). These observations indicate that noxious warmth continue to activate neural circuits within VNC even in the absence of central brain function and thermoreceptor dTRPA1. Taken together our data suggests that cortex glial in the VNC is an important regulator of temperature induced seizures.

## Discussion

In the mammalian nervous system existence of glial heterogeneity across brain regions has been well established (18–20). However, lack of intersectional genetic strategies to label them in a region-specific manner has limited our ability to understand their functional significance (18, 19). As a result, currently only few examples exist that demonstrate functional significance of such glial heterogeneity across mammalian brain regions. For instance, in the mice spinal cord, sensory and motor circuits are spatially segregated into the dorsal and ventral horns respectively. Astrocytes in the ventral horn specifically express axon guidance protein semaphorin3a (Sema3a), conditional deletion of which leads to selective loss of local α-motor neurons, resulting in reduced peak strength behavior in mice (18, 42). Astrocytes in hypothalamic arcuate nucleus are known to interact with pro-opiomelanocortin (POMC) and agouti-related peptide (AgRP) expressing neurons and modulate their synaptic transmission. Astrocyte specific deletion of leptin receptor results in a reduction of synaptic coverage by astrocytic membrane processes and thus specifically affecting POMC and AgRP neuron function leading to leptin resistance, increased fasting or ghrelin-induced hyperphagia in mice (43, 44).

Recently, generation of glial cell specific transcriptional atlas in *Drosophila* suggested that glia exhibit more diversity at the level of morphology rather than at the level of transcription (23). However, existence of transcriptional differences within same glial subtypes cannot be completely ruled out, as sequence depth limits detection of transcripts that are expressed in low levels (23). Existence of *Drosophila* transgenic Gal4 lines that specifically label glial subtypes within same class across brain regions does support possible transcriptional heterogeneity (22).

In the current study, we were able to optimize gene expression in cortex glial subtypes across brain subregions, using Gal4, splitGal4, Gal80, LexA and Killer zipper approaches. Functional specialization of cortex glia across brains regions have not been investigated in *Drosophila*. Here we used seizures in *Drosophila* as a model to study functional significance of the same class of cortex glial population across brain regions including OL, CB and VNC in suppression of seizures. We found that cortex glia in different parts of the brain differentially regulate seizure types including light and temperature sensitive seizures. Our results suggests that CB specific cortex glia are important in suppression of light inducible seizures, whereas VNC specific cortex glia are important in suppression of temperature sensitive seizures. Although, underlying mechanisms are currently unclear, we propose that cortex glia locally regulate neural circuits to prevent hyperactivation, synchronization and spread of neuronal activity across brain regions and thus prevent occurrence of seizures.

*Drosophila* OL and CB constitute neuronal circuits that process visual information (45, 46). We have previously shown that visual transduction *via* norpA mediated signaling is essential for light inducible seizures in *cpes* mutants (27). c*pes* mutants show severe defects in cortex glial morphology in all parts of the brain and the defective cortex glia are responsible for development of light inducible seizures in *cpes* mutants. Further, *cpes* mutants show elevated neuronal activity during seizure episodes *in vivo* (27). In this study we found that CB specific cortex glia are sufficient to suppress light inducible seizures in *cpes* mutants, suggesting important role of neuron-glia interactions in this region. Absence of cortex glia likely affect neuronal organization and function in several ways including regulation of neuroblast proliferation (47–51), proper wiring of neuronal circuits (48, 52), phagocytic clearance of dead neurons (53–57), and maintaining ion balance (28, 29). Thus, it is possible that visual stimulation enhances neuronal activity of CB and lack of cortex glial cell support results in excessive neuronal activity resulting in light inducible seizures.

Glial calcium signaling is involved in regulation of temperature sensitive seizures (29, 31, 58). Acute elevation of cortex glial calcium levels was shown to trigger excessive neuronal activity leading to seizure like activity (31). The *zyd^1^* mutant cortex glia show elevated basal Ca^2+^ levels and calmodulin dependent signaling is critical for temperature sensitive seizures in these mutants (29, 31). Shirley, W. et al., have recently shown that impaired K+ buffering due to enhanced calcineurin dependent endocytosis of K2P leak channel Sandman and other glial membrane proteins is the primary cause for temperature sensitive seizures in *zyd^1^* mutants (29). In this study, we found that Zyd expression in VNC specific cortex glia is essential for suppression of heat inducible seizures in *zyd^1^* mutants. Further, acute influx of Ca^2+^ into VNC specific cortex glia via dTRPA1 expression was sufficient to cause heat inducible seizures. Headless *zyd^1^* mutant flies and UAS-dTRPA1 overexpressing flies showed temperature induced seizures suggesting that VNC specific cortex glia is sufficient for regulation of temperature sensitive seizures. Recent studies have shown that voltage-gated ion channels such as Paralytic (Para), Shaker cognate I (Shal) localize strongly to VNC, and axon initial segments of motor neurons, indicating strong need for glia neuron interactions for rapid conduction of action potentials in these regions (59, 60). Further, it was shown that headless flies expressing dTRPA1 in astrocyte like glia became paralyzed upon shifting temperature to 33°C, suggesting acute Ca^2+^ influx-induced paralysis does not require CB function (58). In *Drosophila* VNC is sufficient to carry out many motor functions and that headless flies survive for several hours and maintain position, walk, groom their front legs, and can even learn to avoid electric shock (61). Thus, it is likely that fly VNC is capable of sensing higher temperatures, and that normal glial cell functions are critical for regulation of neuronal activity in this region.

Taken together, we believe that brain region specific rescue approach taken here to address local role of cortex glia in seizures would spur ongoing research involving localized neuron-glia interactions in broader behavioral outcomes.

## Materials and methods

### Fly stock and husbandry

The following stocks have been obtained from Bloomington *Drosophila* stock center (BDSC), including, VT038983-p65(AD) (BDSC#72320), VT038983-Gal4.DBD (BDSC#72724), R65B12-Gal4 (BDSC#39342), nSyb-Gal80 (BDSC#92154), R54H02-Gal4 (BDSC#45784), UAS-TRPA1 (BDSC#26264), LexAop-rCD2-GFP (BDSC#66544), 13x LexAop-KZip+.3xHA (BDSC#76253), TrpA1 mutant (BDSC#26504), UAS-mCD8GFP (BDSC#5130), UAS-mCherry-NLS (BDSC#38424), Wrapper-nlsLexA::p65 (BDSC#94730), TrpA1[-CD-Gal4] (BDSC#67133), TrpA1[ACD-Gal4] (BDSC#67135), TrpA1-Gal4 (BDSC#65465), TrpA1-Gal4 (BDSC#36922), TrpA1-AB-Gal4 (BDSC#67131), *trpA^1^*[1] (BDSC#26504), Pain-Gal4 (BDSC#27894), c*pes* mutants and UAS CPES were generated in the lab and reported before (27).

Nrv2-p65(AD), and Wrapper-Gal4.DBD are kind gifts from (Coutinho-Budd) (21).

*zyd^1^* mutants, UAS NCKX are kind gifts from (Troy Littleton) (31).

Stocks generated in this study include R54D10-Gal4.DBD, R54D10-nlsLexA.p65, R54D10-Gal80, R9F07-Gal4.DBD, and R9F07-Gal80.

### Cloning and transgenesis

R54D10 and R9F07 enhancers were PCR amplified using primers described at Janelia Flylight project website. These primers were designed to be compatible with gateway cloning technology (Primers Table 3). The PCR product was first cloned into pDONR 221 using BP clonase (Cat#11789020), subsequently cloned into destination vectors pBPZPGal4DBD (Addgene#26233), pBPGal80UW6 (Addgene#26236), and pBPnlsLexA p65UW, (Addgene #26230) using LR clonase reaction as described by manufacturer (Thermofisher Scientific Cat#11791020). Sequence confirmed clones were sent out for embryo injection service at BestGene Inc.

### Immunohistochemistry

Drosophila CNS from larvae, pupae and adults were dissected as described before (https://www.janelia.org/project-team/flylight/protocols and (62). Immunostaining of the dissected CNS was carried out as described previously (27). Briefly, fly brains were dissected in a glass well plate with phosphate buffered saline (1x) and fixed with 4% paraformaldehyde (PFA, EMS Cat#15713-S) in PBS for 1 hour at room temperature (RT). Fixed brains were permeabilized by washing with 1xPBS containing 0.1% Triton x100 (PBST) for 3 times each with 20 min interval at RT (25°C) and blocked with 5% normal goat serum (NGS, Gibco REF#16210-064) in PBST for 1 hour at RT. Subsequently, brains were incubated with primary antibody (10μg/ml in 5% NGS, PBST) for 12 hours/overnight at 4°C and followed by brains were washed with PBST (3 times each with 20 min interval at RT). Brains were then incubated with secondary antibody (10μg/ml in 5% NGS PBST) for 12hours/overnight at 4°C and washed with PBST 3 times each with 20 min interval at RT. Brains were stained with DAPI (1:2000 dilution of 10mg/ml stock, Cat#62247) for 10 min at RT, followed by final wash with PBST for 10 min at RT. In the last step all the buffer was removed by pipetting and a drop of mounting media (Vectashield REF#H1000). The brains were then mounted on a glass slide (VWR Cat#48311-703) with SecureSeal imaging spacer (GBL654008) and covered with cover slip (VWR Cat#48393081/Cat#16004-094) and sealed with clear nail polish (Sally Hansen). The anti-GFP rabbit polyclonal antibody and mCherry monoclonal antibody were from Novus biologicals (NB600-300) and Takara (Cat#632543) respectively.

### Light inducible seizures

The assay for light inducible seizures was performed as described previously (27). Briefly, *cpes* mutants were raised at room temperature (20-22°C) and aged at 25°C with 12 hours light and 12 hours dark cycles for 3-4 weeks. Each food vial contained 20-30 flies, and they were then transferred to new vial every two days until the desired age. The day before the assay flies were allowed to adapt to the dark in a black box (12 hours/overnight). Next day, under infra-red light individual fly vials were arranged on a white cardboard box and videos were captured in real time using Sony cyber shot DSCHX400V camera, 3 seconds under infrared light and 2-5 min under white light (GE, LED, Daylight, 100w/15w, 1600 lumens). Up to 100 flies were tested for each genotype and percent or fraction of flies showing light inducible seizures/photosensitive epilepsy was plotted. We have quantified two different behavioral phenotypes that *cpes* mutant flies exhibited upon exposure to light. Those include, flies fell down lying on their back or side (complete seizure), and flies fell down but remain standing on their legs without any movement for at least 20 seconds (partial seizure). Both behaviors were quantified and plotted.

### Temperature sensitive seizures

The assay for temperature sensitive seizures was performed as described previously (29, 31). Briefly, adult male flies were transferred in groups of 10-20 per vial and aged for 3-5 days. On the day of the assay, flies were transferred to empty vial and allowed to rest for 1-2 hours, subsequently each vial was immersed into preheated transparent water bath maintaining temperature at 38.5°C for 2 minutes for monitoring seizure induction and 2-10 min outside the water bath at RT (22°C) for monitoring seizure recovery. Videos were captured using real time using Sony cyber shot DSCHX400V camera. Total number of flies tested in all assays was always more than 40. To analyze temperature induced seizure behavior, we have quantified only flies that show complete seizures, that is, flies are constantly lying on their back or side with legs twitching. Flies walking up and down on the walls were considered wild type or rescue behavior. Seizure measurements were taken at every 10 second interval and plotted as a fraction of flies showing heat induced seizures as a function of time.

## Acknowledgements

This study was funded by the intramural division of the National Cancer Institute, National Institutes of Health (Division of Health and Human Services)(1ZIABC010331-25). The content of this publication does not necessarily reflect the views or policies of the Department of Health and Human Services, nor does mention of trade names, commercial products, or organizations imply endorsement by the U.S. Government. TAG is supported by the FAU Jupiter Life Science Initiative. KS is supported by FAU Neuroscience Graduate Program. We thank Dr. Troy Littleton for zyd1 and UAS-NCKX flies. We thank Dr. Coutinho-Budd for Nrv2-p65(AD) and Wrapper-Gal4 (DBD) flies.

## Author Contribution

G.K, U.A., & J.K.A conceived the study, G.K designed and performed experiments.

---

**Fig.S1.**
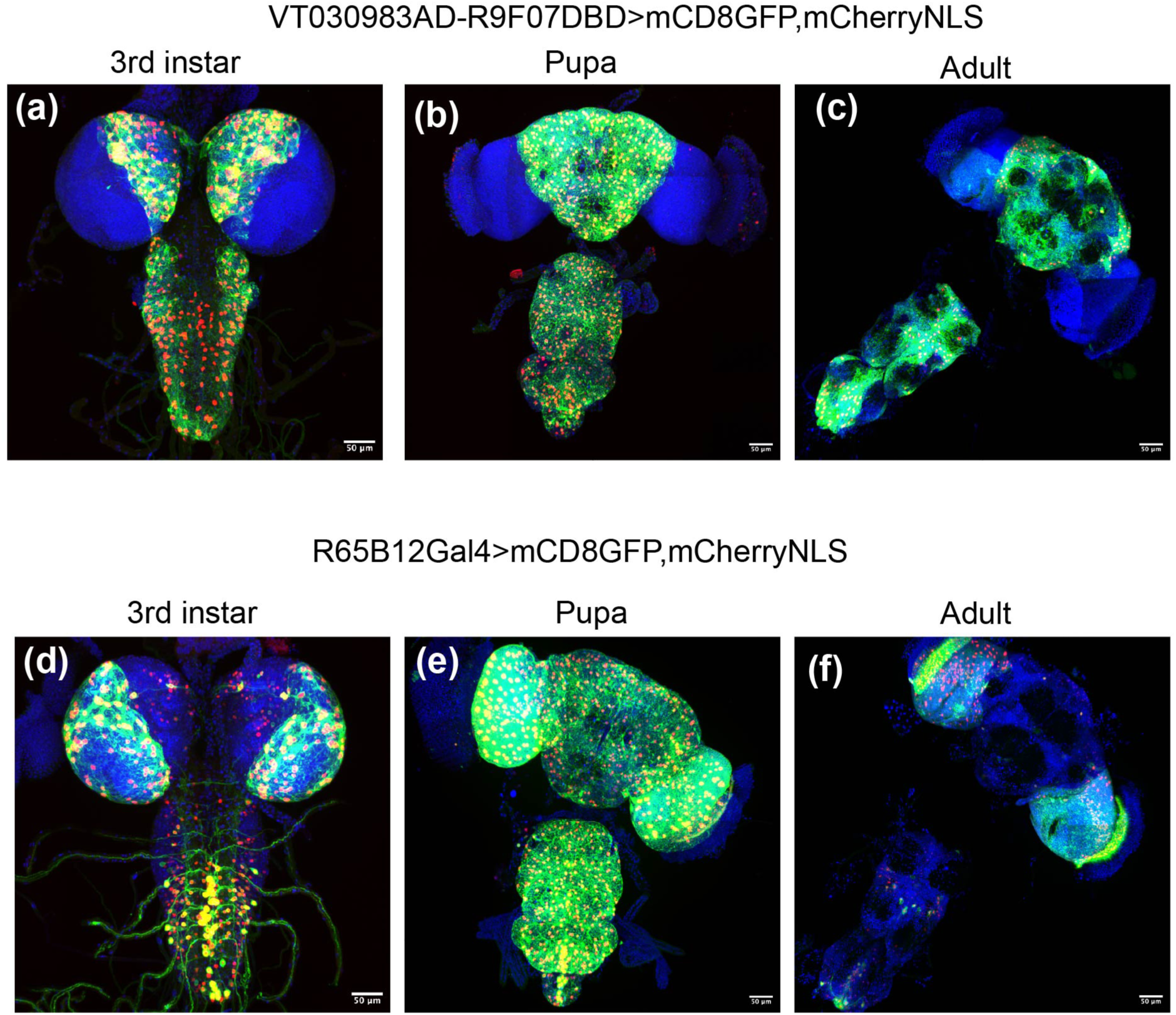
(a-c) Expression pattern of additional CB and VNC specific cortex glial split-Gal4 driver VT030983-p65(AD), R9F07-Gal4.DBD>mCD8GFP, mCherryNLS in 3^rd^ instar CNS (a), 48 hours post pupation CNS and 7-day old adult fly CNS. (d-f) Expression pattern of R65B12-Gal>mCD8GFP, mCherryNLS in 3^rd^ instar CNS, 48 hours post pupation CNS, and 7-day old adult CNS. Scale bar in each of the image corresponds to 50 micrometers.

**Fig.S2.**
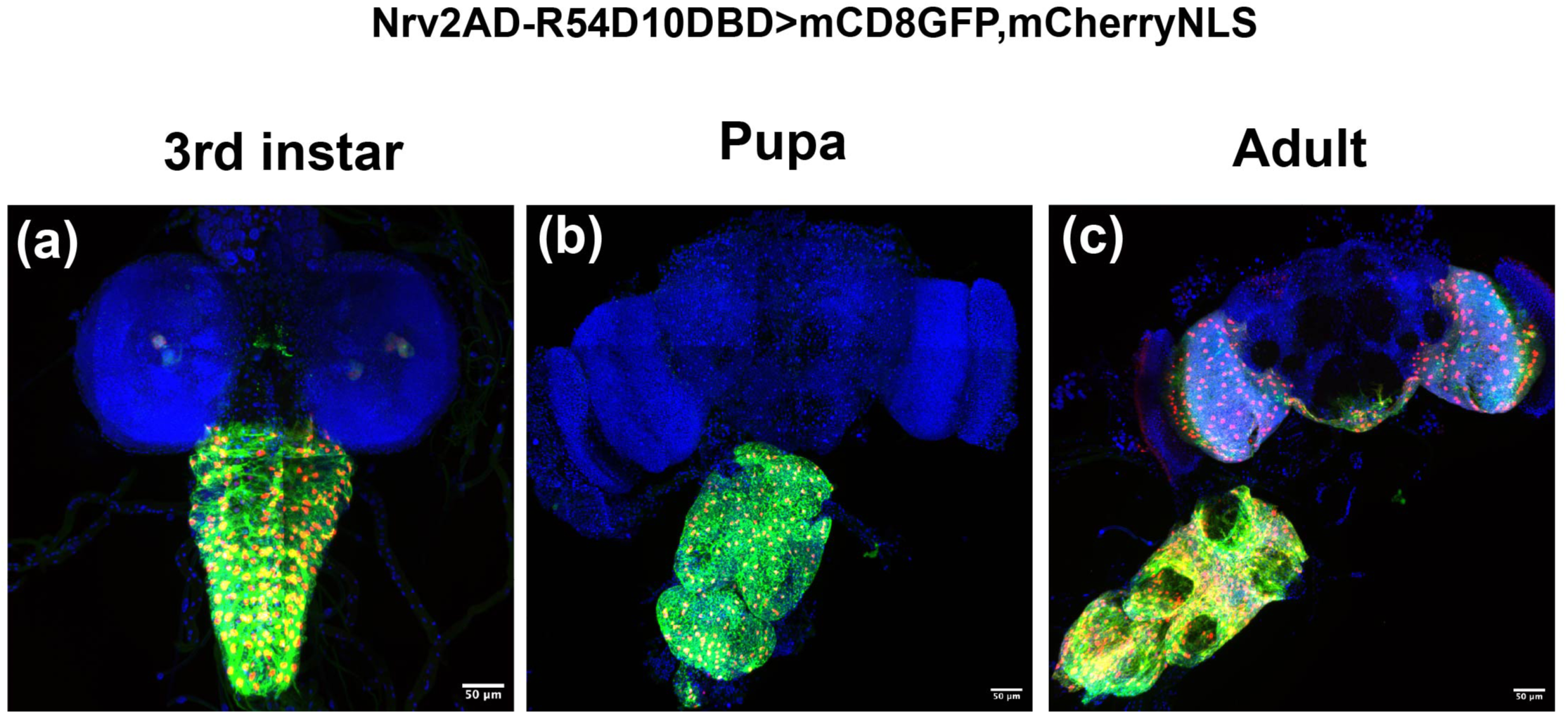
Expression patterns of additional VNC cortex glial specific split Gal4 drivers. (a-c) expression pattern of splitGal4 driver Nrv2-p65(AD), R54D10-Gal4.DBD>mCD8GFP, mCherryNLS in 3^rd^ instar CNS (a), 48 hours post pupation CNS (b), and 7-day old adult CNS (c). Scale bar in each of the image corresponds to 50 micrometers.

**Fig.S3.**
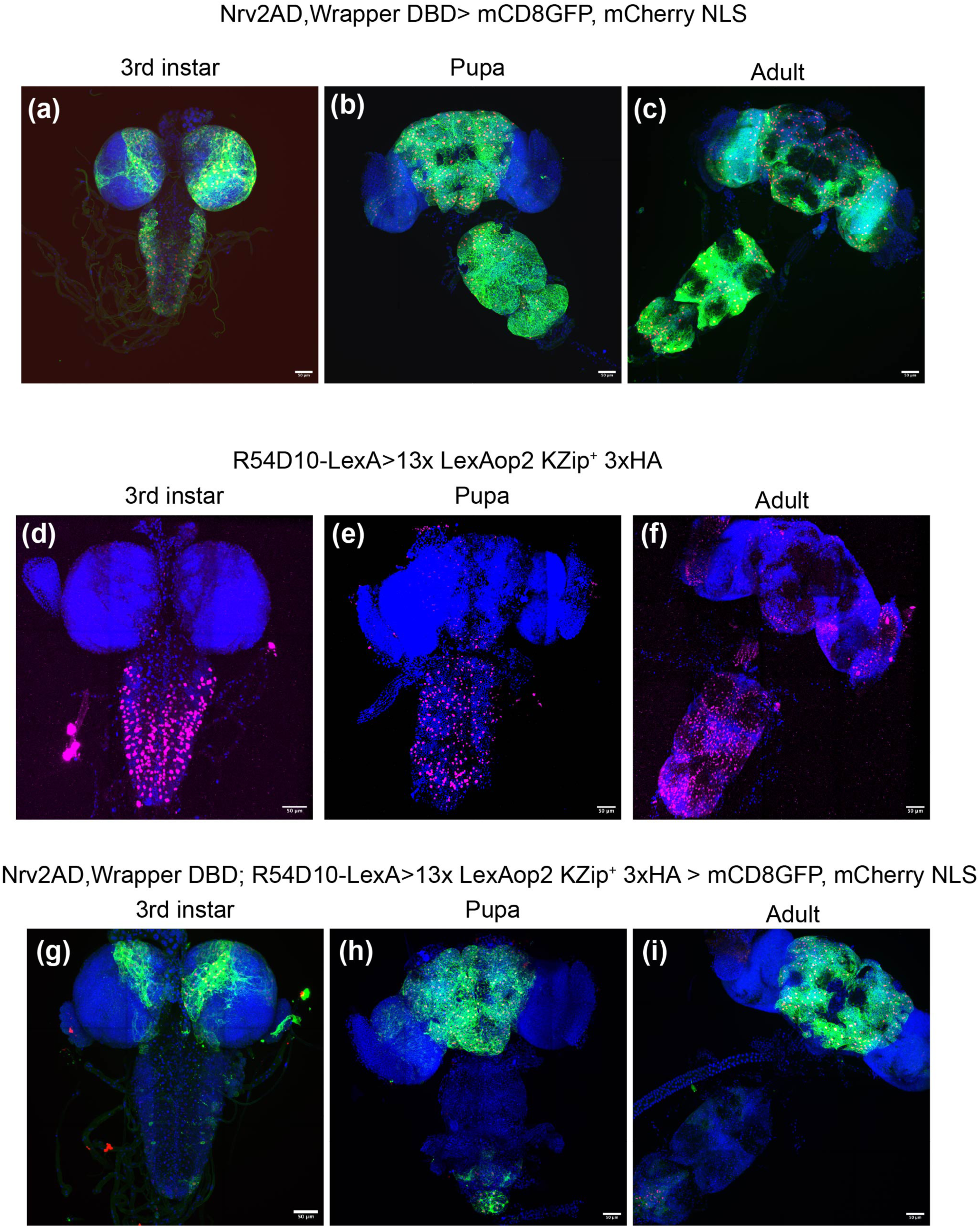
(a-c) Expression pattern of previously described split Gal4 that generically express in all cortex glia, Nrv2-p65(AD), Wrapper-Gal4 DBD (Ctx-Gal4) (21) in 3^rd^ instar CNS (a), 48 hours post pupation CNS (b), and 7-day old adult CNS (c). Ctx-Gal4 drives the expression of mCD8GFP and mCherry NLS. (d-f) expression pattern of R54D10-nlsLexAp65 driving the expression of 13x LexAop2-KZip+. 3xHA. Immunostaining was performed with rabbit anti HA antibody and Alexa Fluor 647 conjugated secondary antibody. 3^rd^ instar CNS (d), 48 hours post pupation CNS (e) and 7-day old adult CNS (f). (g-i) Central brain specific expression pattern was optimized by crossing Ctx-Gal4 with killer zipper system in the VNC using R54D10-nlsLexAp65 and 13xLexAop2-KZip+. The reporters used here are mCD8GFP and mcherry NLS in 3^rd^ instar CNS (g), 48 hours post pupation CNS (h) and 7-day old adult CNS (i). Scale bar in each of the image corresponds to 50 micrometers.

**Fig.S4.**
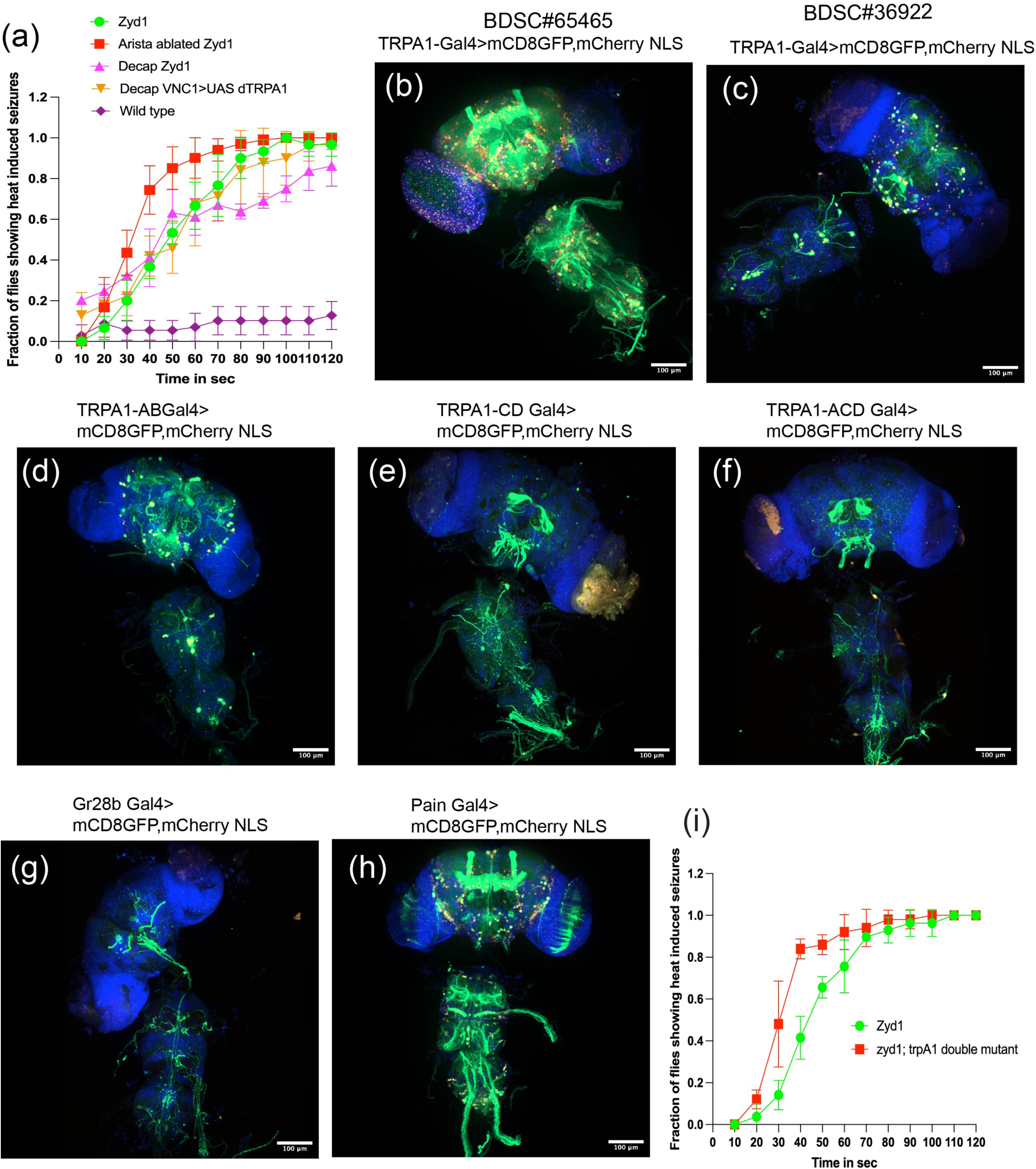
Central brain function is dispensable for initiation of temperature sensitive seizures. (a) Temperature sensitive assay with arista ablated, and decapitated adult flies. Each data point in (a) represents about 50 flies from 3 independent experiments. (b-f) Expression pattern of various TrpA1-Gal4 drivers, (g-h) Expression pattern of Gr28b-Gal4 (g) and Pain-Gal4 (h) drivers in adult CNS. Scale bar in each of the image corresponds to 100 micrometers. (i) measurement of temperature sensitive seizures in *zyd* and *trpA^1^* double mutants. Each data point in (i) represents 50 flies from 5 independent experiments.

**Movie.1. Central brain specific cortex glia plays important role in suppression of light inducible seizures in *cpes* mutants.** Expression of UAS-CPES in VNC specific cortex glia do not suppress light inducible seizures. Whereas expression of UAS CPES in central brain (CB) and optic lobes (OLs) significantly suppressed light inducible seizures. **CB+VNC1**=Nrv2-p65(AD), VT038983-Gal4.DBD>UAS CPES; **OL**=nSybGal80, R9F07Gal80, R65B12-Gal4>UAS CPES; **VNC1**=VT038983-p65(AD), R54D10-Gal4.DBD>UAS CPES; **VNC2**=Nrv2-p65(AD), R54D10-Gal4 DBD> UAS CPES; **CB**=Nrv2-p65(AD), VT038983-Gal4.DBD> UAS CPES, R54D10-nlsLexAp65>13xLexAOP2 Kzip+; ***cpes***=UAS CPES alone. All the above-mentioned rescues are in *cpes* mutant background. All individual movies were assembled and edited into single sequence using Adobe Premiere pro software. Video shown here is the 3x speed of real time.

**Movie.2. Ventral nerve cord specific cortex glia plays important role in suppression of temperature sensitive seizures in *zyd^1^*mutants.** Expression of UAS-NCKX in VNC specific cortex glia is sufficient to suppress temperature sensitive seizures in *zyd^1^* mutants. **CB+VNC**=Nrv2-p65(AD), VT038983-Gal4.DBD>UAS-Zyd; **OL**=R9F07Gal80, R65B12-Gal4> UAS-Zyd; **VNC**=VT038983-p65(AD), R54D10-Gal4.DBD> UAS-Zyd; **CB**=Nrv2-p65(AD), VT038983-Gal4.DBD> UAS-Zyd, R54D10-nlsLexAp65>13xLexAOP2 Kzip+; *zyd^1^*=UAS-Zyd alone. All the above-mentioned rescues are inr *zyd^1^* mutant background. All individual movies were assembled and edited into single sequence using Adobe Premiere pro software. Video shown here is the 3x speed of real time.

**Movie.3. Zyd is required in VNC for the suppression of temperature sensitive seizures in *zyd^1^* mutants.** Expression of Zyd in all cortex glia (All CG) fully suppress temperature sensitive seizures. However, temperature sensitive seizures reoccurred upon repression of Zyd expression in VNC when co-expressed with either R54D10-Gal80 or R9F07-Gal80 in All CG rescue background. *zyd^1^*=UAS-Zyd alone; All CG=R54H02-Gal4>UAS-Zyd; CB+OL=R54H02-Gal4, R54D10-Gal80>UAS-Zyd; OL=R54H02-Gal4, R9F07-Gal80>UAS-Zyd. All the above-mentioned rescues are in *zyd^1^* mutant background. All individual movies were assembled and edited into single sequence using Adobe Premiere pro software. Video shown here is the 3x speed of real time.

**Movie.4. Acute Ca2+ influx into VNC specific cortex glia is sufficient to induce temperature sensitive seizures in wild type flies.** UAS-dTRPA1 is overexpressed in cortex glia in different parts of the brain including VNC, CB and OL. dTRPA1 gets activated only at temperatures above 25°C. Expression and activation dTRPA1 in VNC specific cortex glia is sufficient to induce temperature sensitive seizures in wild type flies. All CG=R54H02-Gal4>UAS-dTRPA1; CB+OL=R54H02-Gal4, R54D10-Gal80>UAS-dTRPA1; OL=R54H02-Gal4, R9F07-Gal80>UAS-dTRPA1; **VNC1**=VT038983-p65(AD), R54D10-Gal4.DBD>UAS-dTRPA1; **VNC2**=Nrv2-p65(AD), R54D10-Gal4 DBD> UAS-dTRPA1. All individual movies were assembled and edited into single sequence using Adobe Premiere pro software. Video shown here is the 3x speed of real time.

**Movie.5. Aristae are dispensable for initiation of temperature sensitive seizures.** Aristae ablated *zyd^1^* mutant flies were subjected to heat shock in a water bath maintaining temperature at 38.5°C for 2 min. Video shown here is the 3x speed of real time.

**Movie.6. Central brain function is dispensable for initiation of temperature sensitive seizures.** *zyd^1^* mutant flies were decapitated under CO_2_ and allowed recovered for 1 hour. Flies were transferred to a Petri dish (60×15mm) and sealed with parafilm. Subsequently, decapitated flies in the dish were immersed in a water bath maintaining temperature at 38.5°C for 2 min to measure temperature sensitivity. **VNC**=VT038983-p65(AD), R54D10-Gal4.DBD>UAS-dTRPA1. Video shown here is the 3x speed of real time.

**Movie.7. Temperature sensitive seizures in *zyd^1^* mutants are independent of thermoreceptor *trpA^1^*function.** Temperature sensitive assay with *zyd^1^* and *trpA^1^*double mutants. Video shown here is the 3x speed of real time.

