## Supplementary material for "Light and temperature sensitive seizures are regulated by spatially distinct cortex glial populations in the central nervous system": Table 1

Table 1: List of Gal4 Drivers from FlyLight project at Janelia whose expression detected in cortex glia of adult brain.

[VT002963](https://flweb.janelia.org/cgi-bin/view_flew_imagery.cgi?line=VT002963)-Trissin receptor

[VT010664](https://flweb.janelia.org/cgi-bin/view_flew_imagery.cgi?line=VT010664)-skywalker

[VT026012](https://flweb.janelia.org/cgi-bin/view_flew_imagery.cgi?line=VT026012)-Ionotrophic receptor64a

[VT038983](https://flweb.janelia.org/cgi-bin/view_flew_imagery.cgi?line=VT038983)-blistry (by) tensin interacts with integrins.

[R9F07](https://flweb.janelia.org/cgi-bin/view_flew_imagery.cgi?line=R9F07)-retn

[R10D01](https://flweb.janelia.org/cgi-bin/view_flew_imagery.cgi?line=R10D01)-bifid (T-box transcription factor)

[R14C04](https://flweb.janelia.org/cgi-bin/view_flew_imagery.cgi?line=R14C04)-nrv2 (non-catalytic component of Na+ K+

[R15A06](https://flweb.janelia.org/cgi-bin/view_flew_imagery.cgi?line=R15A06)-similar (sima-transcriptional regulator of hypoxia)

[R15B11](https://flweb.janelia.org/cgi-bin/view_flew_imagery.cgi?line=R15B11)-Ork1 Open rectifier K+ channel1

[R17F05](https://flweb.janelia.org/cgi-bin/view_flew_imagery.cgi?line=R17F05)-acyl-CoA synthase long chain (AcsI)

[R20B06](https://flweb.janelia.org/cgi-bin/view_flew_imagery.cgi?line=R20B06)-nAChRalpha3

[R24F07](https://flweb.janelia.org/cgi-bin/view_flew_imagery.cgi?line=R24F07)-anachronsm (ana secreted glycoprotein expressed in glia)

[R30E04](https://flweb.janelia.org/cgi-bin/view_flew_imagery.cgi?line=R30E04)- (ocelliless) transcriptional regulator, regulate rhodopsin expression.

[R31F07](https://flweb.janelia.org/cgi-bin/view_flew_imagery.cgi?line=R31F07)-string, tyrosine protein phosphatase

[R33A09](https://flweb.janelia.org/cgi-bin/view_flew_imagery.cgi?line=R33A09)-SNF4/AMP activated protein kinase gamma subunit-lipid metabolism.

[R40B10](https://flweb.janelia.org/cgi-bin/view_flew_imagery.cgi?line=R40B10)-chinmo-BTB-zinc finger transcription factor

[R47A02](https://flweb.janelia.org/cgi-bin/view_flew_imagery.cgi?line=R47A02)-Liprin gamma

[R54H02](https://flweb.janelia.org/cgi-bin/view_flew_imagery.cgi?line=R54H02)-wrapper

[R64C02](https://flweb.janelia.org/cgi-bin/view_flew_imagery.cgi?line=R64C02)-Na pump a subunit

[R64C07](https://flweb.janelia.org/cgi-bin/view_flew_imagery.cgi?line=R64C07) Na pump a subunit

[R73F09](https://flweb.janelia.org/cgi-bin/view_flew_imagery.cgi?line=R73F09)-octa2R-a2-adrenergic like octopamine receptor

[R77A03](https://flweb.janelia.org/cgi-bin/view_flew_imagery.cgi?line=R77A03)-MFS9-Major facilitator superfamily transporter 9

[R81G12](https://flweb.janelia.org/cgi-bin/view_flew_imagery.cgi?line=R81G12)-NFAT-NFAT nuclear factor

[R91G05](https://flweb.janelia.org/cgi-bin/view_flew_imagery.cgi?line=R91G05)-sidestepVIII-expressed in RP neurons and adult head and pCC neurons
