## Supplementary material for "Light and temperature sensitive seizures are regulated by spatially distinct cortex glial populations in the central nervous system": Table 2

**Table 2: List of Gal4/Split-Gal4/LexA/Gal80/Kzip combinations used in this study for brain region specific cortex glial expression.**

| Drivers |  |  |  |  |  |
| --- | --- | --- | --- | --- | --- |
| Brain region | Genotype | 3^rd^ instar | pupa | adult | Comment |
| CB+VNC  (Fig.1a-c) | Nrv2-p65(AD), VT038983-Gal4(DBD). | CB+VNC | CB+VNC | CB+VNC+OL | Additional expression in adult OLs |
| CB+VNC  (Fig.S1a-c) | R9F07-p65(AD), VT038983-Gal4(DBD) | CB+VNC | CB+VNC | Mostly CB+VNC | Weak expression in one of the two OLs |
| OL  (Fig.S1d-f) | R65B12-Gal4 | OL | OL+CB+VNC | OL | Expression in all cortex glia in pupal stage |
| OL  (Fig.1d-f) | R65B12-Gal4, R9F07-Gal80, nSyb-Gal80 | OL | OL | OL | Restricted expression to OLs in all stages |
| VNC  (Fig.S2a-c) | Nrv2-p65(AD), R54D10DBD | VNC | VNC | VNC+SEG+OL | Additional expression in OLs and SEG in adult brain |
| VNC  (Fig.1g-i) | VT038983 (AD)  R54D10.DBD | VNC+CB* | VNC | VNC+SEG+OL | *Additional expression in 3^rd^ instar central brain neuroblast lineages. In the adult brain expression also occurs in SEG and OL |
| CB  (Fig.1j-l) | Nrv2-p65(AD), VT038983-Gal4(DBD), R54D10-(LexA), 13xLexAop2 Kzip+. | CB* | CB* | CB* | *Tip of the VNC continues to show expression in 3^rd^ instar and pupa but weakly in adult. |
| CB  (Fig.S3g-i) | Nrv2-p65 (AD), Wrapper Gal4 (DBD), R54D10-(LexA), 13xLexAop2 Kzip+. | CB* | CB* | CB* | *Tip of the VNC continues to show expression in 3^rd^ instar and pupa but weakly in adult. |
| All cortex glia (Ctx-Gal4)  Fig.S3a-c | Nrv2-p65 (AD), Wrapper Gal4 (DBD), | All cortex glia | *All cortex glia | All cortex glia | *Expression is weaker in optic lobes |
| All cortex glia  (Fig.2a & b) | R54H02-nlsLexAp65, LexAop2-rCD2-GFP | NA | NA | All cortex glia | Expresses in all cortex glia of the adult CNS. |
| VNC+ CB CPES rescue  (Fig.2c) | R54H02-nlsLexAp65, LexAop2-rCD2-GFP, Nrv2-p65(AD) and pVT038983-Gal4.DBD, UAS CPES | NA | NA | Full rescue in VNC+CB but partial rescue in OL | Full rescue in VNC+CB but partial rescue in OL |
| VNC CPES rescue  (Fig.2d) | R54H02-nlsLexAp65, LexAop2-rCD2-GFP, VT038983-p65(AD), and R54D10-Gal4.DBD, UAS CPES | NA | NA | Full rescue in VNC and SEG. No rescue in CB and OLs | Full rescue in adult VNC and SEG. No rescue in CB and OLs. |
| OL CPES rescue  (Fig.2e) | R54H02-nlsLexAp65, LexAop2-rCD2-GFP, R65B12-Gal4, R9F07Gal80 | NA | NA | Full rescue in OLs and partial rescue in CB. | Full rescue in OLs and partial rescue in CB. No rescue in VNC. |
| All cortex glia  (Fig.4a-c) | R54H02-Gal4 (Wrapper Gal4) | All cortex glia | All cortex glia | All cortex glia | All cortex glia |
| CB  (Fig.4d-f) | R54H02-Gal4, R54D10 Gal80 | CB+OL | CB+OL | CB and weakly in OL | Repression is incomplete in adult OLs. |
| OL  (Fig4.g-i) | R54H02 Gal4,  R9F07-Gal80 | *OL | OL | OL | *Weak expression in 3^rd^ instar VNC. |
