## Supplementary material for "Light and temperature sensitive seizures are regulated by spatially distinct cortex glial populations in the central nervous system": Table 3

| R54D10 FW | ggggACAAgTTTgTACAAAAAAgCAggCTTCgTCTACATCgACAAggCATCCgAgT |
| --- | --- |
| R54D10 Rev | ggggACCACTTTgTACAAgAAAgCTgggTCgTTTgCggTgCgATCgCCATTTTTg |
| R9F07 FW | ggggACAAgTTTgTACAAAAAAgCAggCTTCCACCCAgggTgATCCAATTCgC |
| R9F07 Rev | ggggACCACTTTgTACAAgAAAgCTgggTCTCAgTgCCAAATgCCAAA |

Table.3: List of primers used for cloning in this study.
